# Optimization of the cultivation conditions of indigenous wild yeasts and evaluation of their leavening capacity

**DOI:** 10.1101/553818

**Authors:** Elsa Beyene Gebreslassie, Anteneh T. Tefera, Diriba Muleta, Solomon K. Fantaye, Gary M. Wessel

## Abstract

Ethiopia has a high demand for baker’s yeast in the bread and beverage industries. Unfortunately, Ethiopia has no producing plant for baker’s yeast and instead relies on costly imports. The objective of this work was to identify the most productive and useful indigenous baker’s yeasts isolated from local fermented foods and drinks, honey and Molasses using leavening ability as the major metric. Six of the test isolates produced a maximum cell mass at 30°C, pH of 5.5 and 48 hours of incubation. Isolate AAUTf1 did not produce hydrogen sulfide, while isolates AAUTf5, AAUTj15 and AAUSh17 produced low levels of this chemical, and isolates AAUMl20 and AAUWt21 produced high levels of hydrogen sulfide, neglecting their utility in baking. The leavening performance of isolates AAUTf1 (*Candida humilis*) and AAUTf5 (*Kazachstania bulderi*) had the highest dough volume of 131 cm^3^ and 128 cm^3^ respectively in 120 min. Isolates AAUSh17 (*Saccharomyces cerevisiae*) and AAUTj15 (*Saccharomyces cerevisiae*) raised the dough volume of 127 cm^3^ and 125 cm^3^ respectively, at 60 min compared to commercial yeast (117 cm^3^ in 90 min). The study also revealed that mixed cultures of indigenous yeasts had better leavening capacity than single cultures. The co-inoculated cultures of AAUTf1 + AAUTf5 + AAUTj15, AAUTf5 + AAUTj15, and AAUTf1 + AAUTj15 + AAUSh17 reached 143 cm^3^ at 90 min, 141 cm^3^ and 140 cm^3^ both at 60 min, respectively. Thus, the indigenous isolates are candidates for optimizing utilization of yeast for fast promotion and utilization in the bakery industries.

## Introduction

The world population is growing and is expected to reach 9 billion people by the middle of this century [1]. One of the consequences of this increment in population is a higher consumption and a larger demand for processed food such as bread [2]. The greater demand for bread as a staple food for human consumption has led to the development and expansion of the baker’s yeast industry [3].

Bread is a major nutritional component of humans and bread making is one of the oldest processes worldwide, known and practiced for thousands of years [4]. Yeasts are the major microorganism involved in bread making with key role of leavening bread dough Leavening is the metabolic process whereby yeast converts the carbohydrates in the dough to carbon dioxide gas that expands the dough prior to baking [5, 6].

Baker’s yeast (*Saccharomyces cerevisiae*) is the common name for the yeast commonly used as a leavening agent in baking bread and other bakery products, where it converts the fermentable sugars present in the dough into carbon dioxide and ethanol [7]. The fermentative activity of baker’s yeast is essential not only for the rising action of the dough by a production of carbon dioxide but also in a production of the wide range of aromatic compounds identified in bread [8].

Baked foods are widely consumed in Ethiopia and play an important role in the local economy [9]. The bakery sector is constantly growing in Ethiopia due to an increasing demand for bread (particularly commercially prepared bread), constant growth in income, population, urbanization, and due to the shift from traditional consumption habits to fast food. Moreover, a number of alcohol and beverage industries (beer and wine) are active and these industries need tremendous amounts of yeast. As a result, the use of commercial baker’s yeast is increasing day to day in the country.

The supply of commercial yeast in Ethiopia is currently met by importation due to lack of baker’s yeast producing plants in the country [10]. The country spent 293,010,632 ETB (14,650,531.6 US $) in 2016 (CSA 2016) for the imported baker’s yeast. This vital and highly expensive import necessitates alternatives for national development since the raw materials (molasses and wild yeasts) essential to isolate industrial yeasts are locally available.

Many different substrates (fermented foods, fermented beverages, citrus juice, sugarcane juice, molasses and others) are available for the isolation of yeast species [9, 11–13]. However, the leavening capacity of wild yeasts isolated from these substrates *(teff* dough, wheat dough, *shamita, tej,* and molasses) needs proper investigation in order to develop commercial scale production.

Therefore, it is necessary to isolate and develop superior performing baker’s yeast, which would fulfill this demand and thereby save the country enormous expenses. The principal purpose of the present study was to optimize the cultivation conditions of indigenous wild yeasts isolated from local fermented foods and beverages and compared to the commercial baker’s yeast based on their leavening ability in wheat dough.

## Materials and methods

Yeasts isolated from fermented foods (*teff* dough, wheat dough), fermented beverages (*Tej, shamita*) and molasses including the commercial yeast (control) were grown on yeast extract peptone dextrose agar (YEPDA). The isolates were transferred to respective slant medium and preserved at 4°C for further study. The yeast strains used in this study were obtained from my previous research result and were identified using molecular method and the nucleotide sequence was performed at Genwiz, USA. Yeast species name used in this experiment, designation and their source are listed in table 1.

**Table 1.** Yeast species name, designation and their source

| Species name | Designation | Source |
| --- | --- | --- |
| *Candida humilis* (KY102138.1, CBS) | AAUTf | *Teff* dough |
| *Kazachstania bulderi* (KY103628.1, CBS) | AAUTf | *Teff* dough |
| *Saccharomyces cerevisiae* (KY105143.1, CBS) | AAUSh | *Shamita* |
| *Saccharomyces cerevisiae* (KY630581.1, CBS) | AAUTj | *Tej* |
| *Pichia kudriavzevii* (KY104596.1, CBS) | AAUWt | Wheat dough |
| *Pichia fermentans* (KY104550.1, CBS) | AAUMI | Molasses |

### Optimization of cultivation condition for yeast growth

#### Effect of pH on yeasts growth

Isolated yeasts and control (commercial yeast) were separately cultured in yeast extract peptone dextrose (YEPD) broth containing yeast extract 1.0%, peptone 2.0%, and dextrose 2.0%. The pH values were adjusted to 3.5, 4, 4.5, 5 and 5.5 and incubated at 30°C for 48 hours under shaking at 120 rpm [14]. Two 250ml flasks containing 50 ml broth for the listed pH values were each inoculated with 1 ml of a 48 hour-old yeast culture (approximately 1.2 x10^8^ CFU) separately. Optical densities at 600 nm were determined using a spectrophotometer (UV-VIS spectrophotometer, USA) as a measure of growth. The culture medium was used as blank.

#### Effect of temperature on yeast growth

The ability of the isolates including the control to grow at different temperature values was examined by inoculating duplicate flasks with 50 ml YEPD broth medium. The experiment was arranged at four different temperatures values (25, 30, 35, and 40°C) and at optimum pH 5.5 (a result of this study), inoculated with the same number of actively grown yeast cells (48 hours old), 1 ml (approximately 1.2 x10^8^CFU). After 48 hours of incubation optical density were determined the same method as indicated above.

#### Determination of optimum length of time for yeasts growth

The optimum time of incubation for a maximum cell biomass production of each yeast isolate and control (commercial yeast) was determined by incubating cultures at optimum temperature (30°C; result of this study) for 24, 48, 72, 96 and 120 hours. The same number of active yeast cells grown in YEPD for 48 hours, 1 ml (approximately 1.2 x10^8^CFU) was inoculated in duplicates in 50 ml YEPD broth in 250 ml flasks. The best incubation time for growth and maximum biomass production was detected by measuring optical density as indicated above.

#### The interaction effect of temperature, pH and incubation time on yeast growth

The 48 hours old yeast (30°C, 120 rpm) cultured in YEPD broth were inoculated with the same number of actively grown yeast cells 1 ml (approximately 1.2 x10^8^ CFU) at five pH levels (3.5, 4, 4.5, 5 and 5.5) and incubated at 25, 30, 35 and 40°C being shaken at 120 rpm for five days. Samples were taken and analyzed at interval of 24, 48, 72, 96 and 120 hours. Optimum temperature, pH and incubation time for yeast growth and maximum biomass production were determined by using spectrophotometer at 600 nm (UV-VIS spectrophotometer, USA).

### Test of hydrogen sulfide production

To examine production H_2_S (associated with an off-flavor and unpleasant taste), test strains and the control (commercial yeast) were streak cultured on Bismuth Sulfate Agar (BSA) plates and incubated at 30°C for 2 days. Colonies that exhibited significant black color along the line of inoculation on BSA plates indicated hydrogen sulfide production [15]. Positive strains were discarded as their palatability for humans is compromised.

### Preparation of wheat bread with selected yeast isolates

#### Analysis of bread leavening potential of selected yeasts

Bread dough was prepared with candidate isolates to observe the baking potency according to [3]. Selected yeast species and the control for dough making were grown in YEPD broth for 48 hours at optimum temperature of 30°C being shaken at 120 rpm. Samples (10 ml each) were centrifuged for 10 min at 5,000 rpm, washed twice with deionized water, and the supernatant was discarded. The sedimented yeast biomass with moisture was transferred to pre-weighed filter paper, dried overnight at 60°C, and stored in a desiccator until a constant weight was obtained [10]. The yeast culture was harvested and weighed using an analytical balance (FA2104, China).

Prepared dough for this assay contained wheat flour (50 g), harvested yeast culture (0.5 g), table sugar (0.2 g). These ingredients were properly mixed with distilled water (40 ml) and added into 250 ml measuring cylinders. Commercial yeast (Saf-instant, from Turkey) was used separately as a positive control to ferment the dough. Another set of dough formulation that did not contain any yeast sample was prepared as the negative control. The dough samples were left to ferment at ambient (24°C) and 30°C temperatures for 3 hours. The dough volume was determined by measuring the mean of volume increment at every 30 min interval for 3 hours. All dough samples were covered using aluminum foil.

#### Formulation of mixed culture and testing bread leavening potential

The effect of combined (mixed) yeast culture on leavening activity was evaluated. Dough was prepared with commercial yeast and without yeast as positive and negative control. The ingredients used for the dough preparation were wheat flour (50 g), harvested yeast culture (0.5 g), table sugar (0.2 g) and distilled water (40 ml). The ingredients were mixed to homogeneity and incubated at the optimum temperature of 30°C (based on previous result of this study). Single and mixed isolates of yeast cultures used for this test are listed in (Table 2). Two replicates were performed for each type of dough fermentation. The rising power of the combined (mixed) and single (mono) yeast was determined by recording the dough volume increment starting from zero to two hours at 30 min interval. Aluminum foil was used to cover the dough containing measuring cylinders.

**Table 2.** Formulation for bread dough preparation.

| Mixed culture | Harvested yeast culture<br>in gram | Wheat flour in<br>gram | Table sugar in<br>gram | dH<sub>2</sub>O in ml |
| --- | --- | --- | --- | --- |
| X1 | 0.5 | 50 | 0.2 | 40 |
| X2 | 0.5 | 50 | 0.2 | 40 |
| X3 | 0.5 | 50 | 0.2 | 40 |
| X4 | 0.5 | 50 | 0.2 | 40 |
| X5 | 0.5 | 50 | 0.2 | 40 |
| X6 | 0 | 50 | 0.2 | 40 |
| X1+X2 | 0.5 | 50 | 0.2 | 40 |
| X1+X3 | 0.5 | 50 | 0.2 | 40 |
| X1+X4 | 0.5 | 50 | 0.2 | 40 |
| X2+X3 | 0.5 | 50 | 0.2 | 40 |
| X2+X4 | 0.5 | 50 | 0.2 | 40 |
| X3+X4 | 0.5 | 50 | 0.2 | 40 |
| X1+X2+X3 | 0.5 | 50 | 0.2 | 40 |
| X2+X3+X4 | 0.5 | 50 | 0.2 | 40 |
| X1+X3+X4 | 0.5 | 50 | 0.2 | 40 |
| X1+X2+X3+X4 | 0.5 | 50 | 0.2 | 40 |

Note: nomination for isolates X1 (AAUTf1), X2 (AAUTf5), X3 (AAUTj15), X4 (AAUSh17), X5 (+ve control/commercial yeast), X6 (-Ve control)

### Statistical analysis of the experiments

The analysis of variance (ANOVA) of the different sets of experiments or combinations was performed using R software version 3.3.1 [16]. The mean comparison was made using least significant difference (LSD) test at 5% significant level.

## Results

### Optimization of cultivation conditions for yeast growth

#### The effect of pH on yeast biomass

Growth of the isolates varied at different pH values (Table 3). Although all isolates grew at each of the pH levels tested, the minimum and maximum growth yield was observed at pH 3.5 and 5.5 values, respectively. The maximum biomass yields of OD reading at 600 nm reading at pH 5.5 for isolate AAUMl20, AAUSh17, AAUWt21 and AAUTj15 were 2.57, 2.45, 2.25 and 2.23 respectively. Isolate AAUM120 was found to gain the highest biomass yield at the same pH value. However, the maximum biomass yield (1.844) for the control was achieved at pH 5. There were significant (p < 0.05) differences among the biomass yield of the isolates at each pH values (Table 3).

**Table 3.** Mean biomass of Yeasts under different pH ranges

| Isolate | pH3.5 | pH4 | pH4.5 | pH5 | pH5.5 |
| --- | --- | --- | --- | --- | --- |
| AAUTf1 | 0.64<sup>op</sup> | 1.56<sup>defgh</sup> | 1.50<sup>defghi</sup> | 1.82<sup>bcdef</sup> | 1.91<sup>bcde</sup> |
| AAUTf5 | 0.57<sup>p</sup> | 1.43<sup>fghij</sup> | 1.51<sup>defghi</sup> | 1.63<sup>defgh</sup> | 1.85<sup>bcdef</sup> |
| AAUTj15 | 0.76<sup>mnop</sup> | 1.28<sup>ghijk</sup> | 1.2<sup>hijklm</sup> | 1.56<sup>defgh</sup> | 2.23<sup>abc</sup> |
| AAUSh17 | 0.74<sup>nop</sup> | 1.05<sup>jklmno</sup> | 1.2<sup>hijklm</sup> | 1.81<sup>bcdef</sup> | 2.45<sup>a</sup> |
| AAUM120 | 0.83<sup>lmnop</sup> | 1.09<sup>ijklmn</sup> | 1.25<sup>ghijkl</sup> | 1.47<sup>efghij</sup> | 2.57<sup>a</sup> |
| AAUWt21 | 0.73<sup>nop</sup> | 0.97<sup>klmnop</sup> | 1.68<sup>defg</sup> | 1.94<sup>bcd</sup> | 2.25<sup>ab</sup> |
| Control | 1.67<sup>defg</sup> | 1.68<sup>defg</sup> | 1.67<sup>defg</sup> | 1.84<sup>bcdef</sup> | 1.77<sup>cdef</sup> |

Note: CY stands for commercial yeast
Means with the same letter are not significantly different at $p < 0.05$ .

#### The effect of temperature on yeast biomass

The yeast isolates grew at all temperature values (Table 4). The maximum biomass yield for all the six yeast isolates and the control was at 30°C and the minimum biomass yield for all the isolates (including the control) was above 35°C. At 30°C, the AAUMl20 isolate exhibited the maximal growth but biomass yield of all the isolates was significantly higher at 30°C than at all other temperature values (25°C, 35°C and 40°C) (Table 4).

**Table 4.** Mean biomass of potent yeasts under different temperature ranges (values given are O.D.600).

| Isolate | 25°C | 30°C | 35°C | 40°C |
| --- | --- | --- | --- | --- |
| AAUTf1 | 0.6<sup>hijk</sup> | 1.9<sup>d</sup> | 0.56<sup>hijk</sup> | 0.47<sup>jk</sup> |
| AAUTf5 | 0.61<sup>hij</sup> | 1.87<sup>de</sup> | 0.58<sup>hijk</sup> | 0.40<sup>k</sup> |
| AAUTj15 | 0.56<sup>hijk</sup> | 2.21<sup>c</sup> | 0.59<sup>hijk</sup> | 0.46<sup>jk</sup> |
| AAUSh17 | 0.64<sup>hij</sup> | 2.39<sup>ab</sup> | 0.72<sup>h</sup> | 0.52<sup>hijk</sup> |
| AAUMI20 | 0.67<sup>hi</sup> | 2.6<sup>a</sup> | 0.69<sup>hi</sup> | 0.53<sup>hijk</sup> |
| AAUWt21 | 0.51<sup>ijk</sup> | 2.27<sup>bc</sup> | 0.57<sup>hijk</sup> | 0.52<sup>ijk</sup> |
| control | 1.42<sup>f</sup> | 1.79<sup>de</sup> | 1.63<sup>e</sup> | 1.11<sup>g</sup> |

Note: CY stands for commercial yeast
Means with the same letter are not significantly different at $p < 0.05$ .

#### Effect of incubation period on yeast biomass yield

The effect of incubation time on the growth rates of the six isolates and the control at optimum temperature (30°C) and pH of 5.5 is shown in Table 5. Maximum biomass yield was obtained for all the yeast isolates of this study at 48 hours but the minimum biomass yield was recorded decreasing thereafter to the minimum level at 120 hours. Isolate AAUMl20 achieved the highest biomass yield (2.57, OD_600nm_) at 48 hours of incubation time and optimum temperature 30°C followed by isolate AAUSh17 (2.41, OD_600nm_) under the same incubation time and temperature. Table 5 documents the biomass yield for all isolates and we conclude from these results that, except for the control with an optimal incubation time of 72 hours, all other isolates peaked growth characteristics at 48 hours.

**Table 5.** Mean growth of Yeasts under different incubation time ranges (OD_600nm_)

| Isolate | 24 hours | 48 hours | 72 hours | 96 hours | 120 hours |
| --- | --- | --- | --- | --- | --- |
| AAUTf1 | 1.6<sup>gh</sup> | 1.96 | 1.05<sup>jk</sup> | 0.59<sup>o</sup> | 0.36<sup>q</sup> |
| AAUTf5 | 1.49<sup>h</sup> | 1.81<sup>ef</sup> | 0.79<sup>n</sup> | 0.94<sup>klm</sup> | 0.52<sup>op</sup> |
| AAUTj15 | 1.68<sup>efg</sup> | 2.12<sup>cd</sup> | 1.33<sup>i</sup> | 0.84<sup>mn</sup> | 0.61<sup>o</sup> |
| AAUSh17 | 1.79<sup>ef</sup> | 2.43<sup>b</sup> | 0.91<sup>lmn</sup> | 0.81<sup>mn</sup> | 0.43<sup>pq</sup> |
| AAUMI20 | 1.7<sup>efg</sup> | 2.6<sup>a</sup> | 0.99<sup>kl</sup> | 0.83<sup>mn</sup> | 0.43<sup>pq</sup> |
| AAUWt21 | 1.66<sup>fg</sup> | 2.21<sup>c</sup> | 1.15<sup>j</sup> | 0.98<sup>kl</sup> | 0.53<sup>op</sup> |
| CY(Control) | 1.31<sup>i</sup> | 1.74<sup>efg</sup> | 1.99<sup>cd</sup> | 1.81<sup>e</sup> | 1.8<sup>e</sup> |

Note: CY stands for commercial yeast
Means with the same letter are not significantly different at $p < 0.05$ .

#### Combined effect of temperature, pH and incubation time on yeast biomass yield

The maximum cell density 1.89, 1.82, 2.17, 2.41, 2.56 and 2.23 of OD at 600nm for isolates AAUTf1, AAUTf5, AAUTj15, AAUSh17, AAUMl20 and AAUWt21 respectively, were obtained at 30°C, pH 5.5 and 48 hours of incubation (data not shown). On the other hand, the maximum biomass yield for control yeast (2.0, OD_600nm_) was achieved when the temperature, pH and incubation time was at 30°C, 5 and 72 hours, respectively. We observed a significant difference (p< 0.05) among the treatments on the combined effect of temperature, pH and incubation time with regard to biomass yield. The minimum biomass yield was measured for the isolates AAUTf1 (0.21), AAUTf5 (0.25), AAUTj15 (0.27), AAUSh17 (0.28), AAUMl20 (0.35), AAUWt21 (0.28) and control (0.49, OD_600nm_) at 40°C, 3.5 pH and 120 hours of incubation time.

### Hydrogen sulfide production by yeast isolates

On the basis of their H_2_S production (Fig 1), the isolates were grouped into three categories (non-producers, low level and high level of H_2_S producers). Accordingly, isolate AAUTf1 did not produce hydrogen sulfide (Fig 1, A), while AAUTf5, AAUTj15 and AAUSh17 produced low levels of hydrogen sulfide. The commercial yeast also produces low levels H_2_S as well (Fig 1, B). Isolates AAUMl20 and AAUWt21 produced high level of hydrogen sulfide (Fig 1, C). Therefore, AAUTf1, AAUTf5, AAUTj15 and AAUSh17 were subjected for further test.

**Fig 1.**
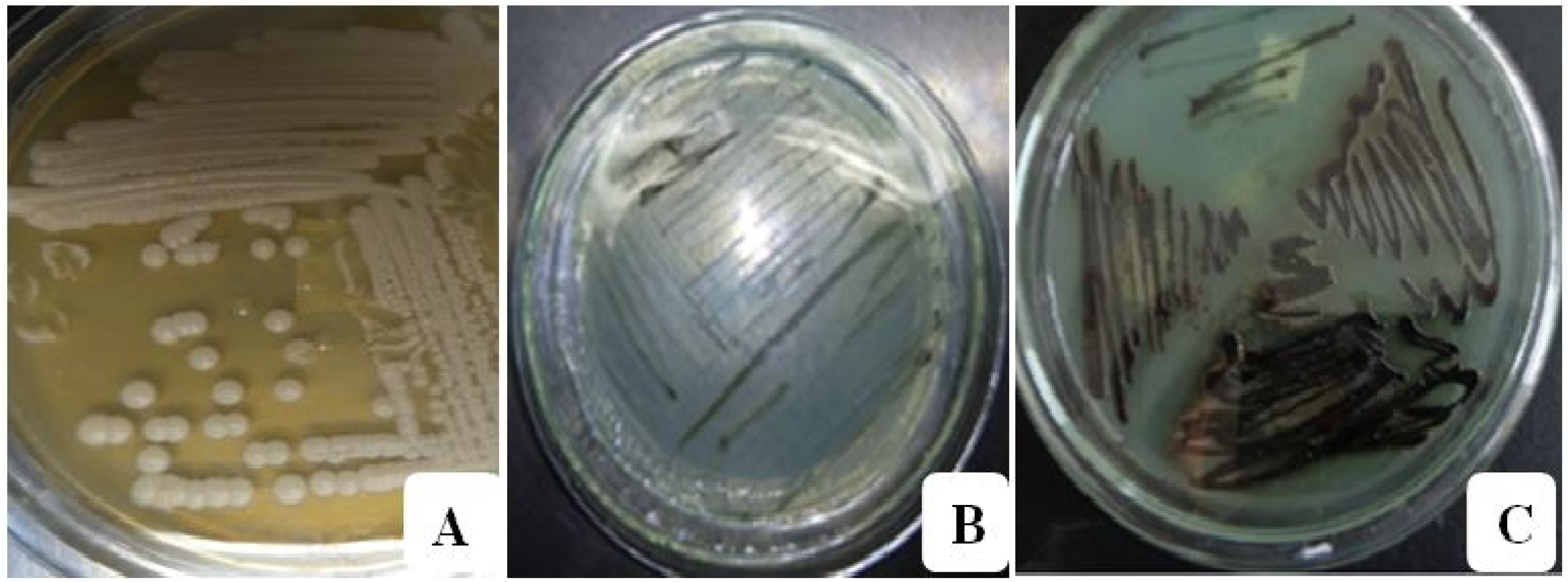
Hydrogen sulfide (H_2_S) gas production by isolates as detected by black readout on Bismuth Sulphate Agar plates. A (AAUTf1) - non producer; B (AAUTf5, AAUTj15, AAUSh17 and Commercial yeast) - low level and C (AAUMl20 and AAUWt21) – high level

### Leavening capacity of isolated yeast strains

The leavening capacity of the non-hydrogen sulphide producer, *C. humilis s*train (AAUTf1), and the low H_2_S producers *K. bulderi* strain (AAUTf5), *S. cerevisiae* strain (AAUTj15), and *S. cerevisiae* strain (AAUSh17) were compared to the commercial *S. cerevisiae*. The results showed that the period of bread dough fermentation at 30°C was short (2 hours) compared to ambient temperature (Table 6). The maximum mean of leavening activity was seen by isolate AAUTf1 (131 cm^3^) at 120 min, which was followed by AAUTf5 (128 cm^3^) at 120 min at 30°C. Similarly, isolates AAUSh17 (127 cm^3^) and AAUTj15 (125 cm^3^) achieved high leavening activity at 60 min at the same temperature, which was not significantly different (p>0.05) with the above isolates (AAUTf1 and AAUTf5). The commercial yeasts had 117 cm^3^ mean rising capacity at 90 min which is lower, and takes longer (p<0.05) than that of the indigenous isolates. Dough left to ferment without yeast (negative control) did not show volume increment within 3 hours of dough fermentation (Table 6).

**Table 6.** Leavening activity of yeast strains at 24°C and 30°C temperature

| Isolates | Temp | Mean of rising dough volume (cm<sup>3</sup>)/ Time (min) |  |  |  |  |  |  |
| --- | --- | --- | --- | --- | --- | --- | --- | --- |
|  |  | 0 min | 30 min | 60 min | 90 min | 120 min | 150 min | 180 min |
| AAUTf1 | 24 °C | 0<sup>i</sup> | 12<sup>hi</sup> | 26<sup>g-i</sup> | 50<sup>d-i</sup> | 63<sup>a-g</sup> | 70<sup>a-g</sup> | 109<sup>ab</sup> |
|  | 30 °C | 0<sup>l</sup> | 32<sup>kl</sup> | 46<sup>i-k</sup> | 73<sup>e-j</sup> | 131<sup>a</sup> | 110<sup>a-e</sup> | 84<sup>c-h</sup> |
| AAUTf5 | 24 °C | 0<sup>i</sup> | 35<sup>f-i</sup> | 51<sup>d-h</sup> | 75<sup>a-g</sup> | 97<sup>a-d</sup> | 81<sup>a-f</sup> | 70<sup>a-g</sup> |
|  | 30 °C | 0<sup>l</sup> | 40<sup>jk</sup> | 80<sup>d-i</sup> | 111<sup>b-d</sup> | 128<sup>a</sup> | 124<sup>ab</sup> | 81<sup>d-i</sup> |
| AAUTj15 | 24 °C | 0<sup>i</sup> | 60<sup>b-h</sup> | 113<sup>a</sup> | 116<sup>a</sup> | 87<sup>a-e</sup> | 73<sup>a-g</sup> | 67<sup>a-g</sup> |
|  | 30 °C | 0<sup>l</sup> | 49<sup>g-k</sup> | 125<sup>ab</sup> | 110<sup>b-d</sup> | 81<sup>d-i</sup> | 84<sup>c-h</sup> | 77<sup>d-j</sup> |
| AAUSh17 | 24 °C | 0<sup>i</sup> | 51<sup>d-h</sup> | 55<sup>c-h</sup> | 59<sup>b-h</sup> | 98<sup>a-d</sup> | 86<sup>a-f</sup> | 82<sup>a-f</sup> |
|  | 30 °C | 0<sup>l</sup> | 48<sup>h-k</sup> | 127<sup>a</sup> | 103<sup>b-e</sup> | 78<sup>d-i</sup> | 86<sup>b-g</sup> | 63<sup>f-k</sup> |
| CY | 24 °C | 0<sup>i</sup> | 39<sup>e-i</sup> | 90<sup>a-d</sup> | 100<sup>a-d</sup> | 103<sup>a-c</sup> | 80<sup>a-f</sup> | 73<sup>a-g</sup> |
|  | 30 °C | 0<sup>l</sup> | 47<sup>h-k</sup> | 101<sup>b-e</sup> | 117<sup>b-c</sup> | 103<sup>b-e</sup> | 95<sup>b-f</sup> | 78<sup>d-i</sup> |
| NC | 24 °C | 0<sup>i</sup> | 0<sup>i</sup> | 0<sup>i</sup> | 0<sup>i</sup> | 0<sup>i</sup> | 0<sup>i</sup> | 0<sup>i</sup> |
|  | 30 °C | 0<sup>l</sup> | 0<sup>l</sup> | 0<sup>l</sup> | 0<sup>l</sup> | 0<sup>l</sup> | 0<sup>l</sup> | 0<sup>l</sup> |

Note: CY- commercial yeast; NC – negative control.
Means with the same letter are not significantly different at $p < 0.05$ .

### Effect of mixed yeast cultures on leavening activity

The combined ability of the four selected yeast isolates AAUTf1 (*C. humilis*), AAUTf5 (*K. bulderi*), AAUTj15 (*S.cerevisiae*) and AAUSh17 (*S.cerevisiae*) on bread dough leavening was tested for additive properties of the yeast. Co-inoculated isolates were compared for their leavening effect to each of the separate isolates and to that of the control, commercial yeast. Results of the three co-inoculated isolates (AAUTf1+ AAUTf5 + AAUTj15) were found highest (143 cm^3^) at 90 min; while the raising volume of dough of as result of co-inoculation of different combination of two (AAUTf5 + AAUTj15) and three (AAUTf1 _+_ AAUTj15 _+_ AAUSh17) yeast isolates was as high as 141 and 140 cm^3^ respectively at 60 min (Table 7). The aroma of the dough prepared using combined isolates was judged better than of the dough prepared by single isolates and the commercial bakery yeast, though admittedly, this is a subjective measurement (data not included).

**Table 7.** Leavening activity of mixed and pure isolates

| Isolate/S | Mean of rising dough volume (cm<sup>3</sup>) at time (min) |  |  |  |  |
| --- | --- | --- | --- | --- | --- |
|  | 0 min | 30 min | 60 min | 90 min | 120 min |
| X1 | 0<sup>B</sup> | 32<sup>zA</sup> | 46<sup>w-A</sup> | 73<sup>n-w</sup> | 131<sup>a-c</sup> |
| X2 | 0<sup>B</sup> | 40<sup>y-A</sup> | 80<sup>l-t</sup> | 111<sup>c-j</sup> | 128<sup>a-d</sup> |
| X3 | 0<sup>B</sup> | 49<sup>v-z</sup> | 125<sup>a-e</sup> | 87<sup>h-t</sup> | 81<sup>k-t</sup> |
| X4 | 0<sup>B</sup> | 48<sup>w-z</sup> | 127<sup>a-d</sup> | 93<sup>g-r</sup> | 78<sup>l-u</sup> |
| X5 | 0<sup>B</sup> | 47<sup>w-A</sup> | 101<sup>d-n</sup> | 117<sup>b-h</sup> | 103<sup>c-m</sup> |
| X6 | 0<sup>B</sup> | 0<sup>B</sup> | 0<sup>B</sup> | 0<sup>B</sup> | 0<sup>B</sup> |
| X1X2 | 0<sup>B</sup> | 49<sup>v-z</sup> | 102<sup>d-m</sup> | 68<sup>q-y</sup> | 68<sup>q-y</sup> |
| X1X3 | 0<sup>B</sup> | 44<sup>x-A</sup> | 98<sup>e-o</sup> | 94<sup>g-r</sup> | 83<sup>j-t</sup> |
| X1X4 | 0<sup>B</sup> | 19<sup>AB</sup> | 96<sup>f-q</sup> | 85<sup>j-t</sup> | 66<sup>r-y</sup> |
| X2X3 | 0<sup>B</sup> | 100<sup>d-n</sup> | 141<sup>ab</sup> | 121<sup>a-h</sup> | 86<sup>i-t</sup> |
| X2X4 | 0<sup>B</sup> | 33<sup>zA</sup> | 124<sup>a-f</sup> | 109<sup>c-k</sup> | 63<sup>s-y</sup> |
| X3X4 | 0<sup>B</sup> | 64<sup>s-y</sup> | 114<sup>b-i</sup> | 77<sup>m-v</sup> | 69<sup>p-x</sup> |
| X1X2X3 | 0<sup>B</sup> | 19<sup>AB</sup> | 97<sup>e-p</sup> | 143<sup>a</sup> | 98<sup>e-o</sup> |
| X2X3X4 | 0<sup>B</sup> | 50<sup>u-z</sup> | 128<sup>a-d</sup> | 93<sup>g-r</sup> | 71<sup>0-x</sup> |
| X1X3X4 | 0<sup>B</sup> | 59<sup>t-z</sup> | 140<sup>ab</sup> | 106<sup>c-l</sup> | 89<sup>h-s</sup> |
| X1X2X3X4 | 0<sup>B</sup> | 51<sup>u-z</sup> | 125<sup>a-e</sup> | 109<sup>c-k</sup> | 89<sup>h-s</sup> |

Note: nomination for isolates X1-AAUTf1; X2-AAUTf5; X3-AAUTj15; X4-AAUSh17; X5-CY (Positive control); X6- Negative control.
Means with the same letter are not significantly different at $p < 0.05$ .

## Discussion

The metabolic and production efficiency of cells depends on many factors such as temperature, pH, incubation period, inoculums size, genetic background [17]. All the isolates, *Candida humilis* (AAUTf1)*, Kazachitania bulderi* (AAUTf5), *Saccharomyces cerevisiae* (AAUTj15 and AAUSh17)*, Pichia fermentans* (AAUMl20) and *Pichia kudrvizivi* (AAUWt21) showed higher biomass at pH of 5.5, temperature of 30°C and incubation time of 48 hours, while the commercial yeast (control) had less biomass. This result shows that the isolated yeasts (this study) had shorter growth times than that of the commercial yeast strain. Similar to this result, [18] has found that yeasts grew maximally at pH 5 to 5.5, 30°C temperature and 72 hours of incubation period.

All the yeast species and strains in this study could tolerate a temperature up to 40°C including the control (Table 4). The ability of yeast to tolerate high temperature suggests that the isolates can withstand excess heat associated with fermentation process and therefore can be used to accomplish fermentation at a wide range of temperature condition. In agreement with this study, [19, 20] have also reported that yeasts can grow at elevated temperatures of 40°C, but the optimal temperature is approximately 30°C.

In the current study, a maximum biomass was obtained at 48 hours of incubation period but the biomass decreased with increasing incubation time. This is supported by the scientific fact that the stationary phase of yeast growth is a period of no growth, when metabolism slows and cell division is stopped due to nutrient deprivation, toxic metabolites and high temperatures which led cells to die and autolyse. In contrary to the present study, [21] have stated that the highest biomass was recorded after 144 hours of incubation period. The difference in these results may be due to the genetic constituent of their cells and cultivation conditions.

The current study has indicated that isolate AAUTf1 did not produce hydrogen sulfide, while AAUTf5, AAUTj15 and AAUSh17 including the commercial yeast produced lower content of this undesirable gas and yet other isolates produced intense dark color on Bismith Sulfate Agar (BSA) medium [15]. Other scholar, [22] also reported that the highly darkened color in Lead Acetate Agar (LAA) indicates a greater amount of hydrogen sulfide production. Therefore, some of the wild yeast isolates in the present study could be a potential candidate for wheat dough leavening for bread making since they showed low production of H_2_S and also had better fermentation ability than the commercial yeast. Furthermore, [23] have demonstrated that yeast strains isolated from fruits and plant parts showed better leavening performance compared to commercial strains.

The results of the present study indicated that the ability of the potent yeast isolates is comparable or even better than the commercial yeast in leavening of bread dough. Similarly, [24] have indicated that yeast strains isolated from fruits showed higher leavening activity than that of the commercial yeast strain. The dough rising power of different brands of baker’s yeasts (from Turkey, China, UK, and Egypt) sold in Egypt have compared and all the yeast strains had maximum leavening activity after 2 hours of fermentation [3], but the highest leavening activity showed by the potent yeast isolates between 1 to 2 hours in the current study. This reveals that the leavening activity of indigenous yeast isolates showed shorter time of fermentation than that of the commercial baker’s yeast making the potent yeast isolates of this study a potential candidate to be developed into commercial bakery yeast strains after further necessary tests.

A combination of the three isolates (AAUTf1 + AAUTf5 + AAUTj15) produced the highest leavening activity compared to single inoculations. Better performance of combined wild yeast isolates (this study) could be due to synergetic contribution of the isolates to the dough leavening action as demonstrated by several investigators [25–30], who reported that a combination of yeasts (non *Saccharomyces cerevisiae + Saccharomyces cerevisiae*) is important for quality bread leavening and baking purpose. Both isolates of AAUTf1 (*Candida humilis*) and AAUTf5 (*Kazachistania bulderi*) of this study are uncommon types of yeasts in baking industries, but they have good leavening ability and aroma than of the commercial yeast (*S.cerevisiae*). Emphasizing the importance of uncommon yeast strains, [31] have demonstrated that many uncommon (non-conventional) types of yeasts are used in baking industries that have the ability to produce unique aroma compounds that *S*. *cerevisiae* lacks.

Overall, it was noticed that the combinations (AAUTf5 + AAUTj15) and (AAUTf1 + AAUTj15 + AAUSh17) of indigenous yeasts isolated from local substrates showed the highest leavening ability of bread indicating the possibility of developing indigenous baker’s yeasts for large scale production. Thus, this can potentially increase the varieties of yeasts and ultimately decrease their importation at huge amount of foreign currencies.

Furthermore, this study may even lead to eventual screening of more indigenous potent yeast blends for local consumption and beyond after conducting various qualifying tests.

## Conclusions

The results of our study demonstrated that fermented foods and drinks harbor potent baker’s yeasts which can be used as dough leavening agents. The optimum growth conditions for yeasts are 30°C temperature, 5.5 pH and 48 hours of incubation. The yeast isolate *Saccharomyces cerevisiae* exhibited good leavening activity and *Candida humilis* and *Kazchistania bulderi* (strains not used before for leavening bread dough) have better capacity of leavening and is concluded to be the most active yeasts to ferment bread dough compared to other strains including commercial yeast strain. Combinations of isolates (mixed culture) with *Saccharomyces cerevisiae* showed higher capacity of wheat dough leavening than the indigenous single isolates (monoculture) and of commercial yeast. Thus, the indigenous isolates are potential candidates that need fast promotion and utilization in bakery industries. Based on the findings of this study it is recommended that further investigation should be undertaken on organoleptic properties and other baker’s yeasts qualifying parameters in order to enhance their desirability and efficiency of the screened strains.

## Acknowledgments

The authors would like to thank Department of Microbial Cellular and Molecular Biology, Addis Ababa University, Molecular Research Center Advanced Laboratory, Addis Ababa, Ethiopia and other members of Brown University, USA for their kind collaboration in providing laboratory and DNA sequencing facilities. We are also thankful of the Ethiopian Ministry of Science and Technology for the financial support to conduct the research work herein.

## References

1. Jensen S, Skibsted LH, Kidmose U, Thybo AK. Addition of cassava flours in bread-making: sensory and textural evaluation. LWT-Food Sci Technol. 2015; 60(1):292–299.

2. Godfray HCJ, Beddington JR, Crute IR, Haddad L, Lawrence D, Muir JF. Food security: the challenge of feeding 9 billion people. Sci. 2010; 327: 812–818.

3. Zaky AS, Nasr NF. Easy and efficient method for measuring the rising power of baker’s yeast. International Food Congress-Novel Approaches in Food Industry 2011; 650–654.

4. Plessas S, Pherson L, Bekatorou A, Nigam P, Koutinas AA. Bread making using kefir grains as baker’s yeast. Food chemis. 2005; 93:585–589.

5. Donalies UE, Nguyen HTT, Stahl U, Nevoigt E. Improvement of Saccharomyces yeast strains used in brewing, wine making and baking. Adv Biochem Eng Biotechnol. 2008; 111:67–98.

6. Edwards WP. The Science of Bakery Products. Royal Society of Chemistry, Cambridge; 2007.

7. Hamelman J. Bread: A Baker’s Book of Techniques and Recipes. John Wiley, New York; 2004.

8. Birch AN, Petersen MA, Hansen AS. The aroma profile of wheat bread crumb influenced by yeast concentration and fermentation temperature. Food Sci Technol. 2013; 50: 480–488.

9. Ashenafi M. The microbiology of Ethiopian foods and beverages: A review. Ethiop J Biol Sci. 2006; 5: 189–245.

10. Tamene M, Dawit A. Evaluation of Yeast Biomass Production using Molasses and Supplements. LAP LAMBERT Academic Publishing, Germany; 2014.

11. Arias CR, Burns JK, Friedrich LM, Goodrich RM, Parish ME. Yeast species associated with orange juice: Evaluation of different identification methods. Appl Env Microbiol. 2002; 68: 1955–1961.

12. Ceccato-Antonini SR, Tosta CD, Silva AC. Determination of yeast killer activity in fermenting sugarcane juice using selected ethanol making strains. Brazil Arch Boil Technol. 2004; 47: 17–39.

13. Aslankoohi E, Herrera-Malaver B, Rezaei MN, Steensels J, Courtin CM, Verstrepen KJ. Non-conventional yeast strains increase the aroma complexity of bread. PloS One. 2016; 11: e0165126.

14. Qureshi SK, Masud T, Sammi S. Isolation and taxonomic characterization of yeast strains on the basis of maltose utilization capacity for bread making. Int J Agri Biol. 2007; 9: 110–113.

15. Jiranek V, Langridge P, Henschke PA. Validation of bismuth containing indicator media for predicting H_2_S producing potential of *Saccharomyces cerevisiae* wine yeast under enological conditions. Am J Enol Vitic.1995; 46: 269–273.

16. Team RC. R: A Language and Environment for Statistical Computing. R Foundation for statistical Computing, Vienna, Austria. 2016, URL https://www.R-project.org/.

17. Supanwong K, Kazuyoshi O, Shinsaku H. Environmental effects on ethanol tolerance of *Zymomonas mobilis*. J. Microbial. Utilization Renewable Resource. 1983; 3: 254–260.

18. Dechassa T, Dawit A. Isolation and selection of ethanol tolerant yeasts for the production of ethanol. M.Sc. Thesis, Addis Ababa University. 2010. Available: http://etd.aau.edu.et/handle/123456789/3625.

19. Cho IH, Peterson DG Chemistry of bread aroma: a review. Food Sci Biotechnol. 2010; 19: 575–582.

20. Nitayavardhana S, Shrestha P, Rasmussen ML, Lamsal BP, van Leeuwen JH, Khanal SK. Ultrasound improved ethanol fermentation from cassava chips in cassava-based ethanol plants. Bioresource technol. 2010; 101: 2741–2747.

21. Mamun-Or-Rashid AN, Dash BK, Chowdhury MN, Waheed MF, Pramanik MK. Exploration of potential baker’s yeast from sugarcane juice: optimization and evaluation. Pak J Biol Sci. 2013; 16: 617–623.

22. Fellers CR, Shostrom OE, Clark ED. Hydrogen sulfide determination in bacterial cultures and in certain canned foods. J bacterial. 1924; 9: 235–249.

23. Noroul A, Ma’aruf Z, Sahilah AG, Mohd AM, Khan A, Wan Aida WM. A new source of *Saccharomyces cerevisiae* as a leavening agent in bread making. Int Food Res J. 2013; 20: 967–973.

24. Ma’aruf AG, Noroul AZ, Sakilah AM, Mohd KA. Leavening ability of yeast isolated from different local fruits in bakery product. Sains Malaysiana. 2011; 40: 1413–1419.

25. Clemente-Jimenez JM, Mingorance-Cazorla L, Martinez-Rodriguez S, Heras-Vazquez FJL, Rodriguez-Vico F. Influence of sequential yeast mixtures on wine fermentation. Int J Food Microbiol. 2005; 98: 301–308.

26. Moreira N, Mendes F, de Pinho RG, Hogg T, Vasconcelos I. Heavy sulfur compounds, higher alcohols and esters production profile of Hanseniaspora uvarum and Hanseniaspora guilliermondii grown as pure and mixed cultures in grape must. Int J Food Microbiol. 2008; 124: 231–8.

27. Domizio P, Romani C, Lencioni L, Comitini F, Gobbi M, Mannazzu I. Outlining a future for non *Saccharomyces* yeasts: Selection of putative spoilage wine strains to be used in association with *Saccharomyces cerevisiae* for grape juice fermentation. Int J Food Microbiol. 2011; 147: 170–80.

28. Crafack M, Mikkelsen MB, Saerens S, Knudsen M, Blennow A, Lowor S. Influencing cocoa flavour using Pichia kluyveri and Kluyveromyces marxianus in a defined mixed starter culture for cocoa fermentation. Int J Food Microbiol. 2013; 167: 103–116.

29. Saerens SM, Swiegers H, Reynolds D. Increasing the sensorial enrichment of white wine with non-*Saccharomyces* yeast strains. Australian and New Zealand Grapegrower and Winemaker. 2013; 599: 94.

30. Steensels J, Snoek T, Meersman E, Nicolino MP, Voordeckers K, Verstrepen KJ. Improving industrial yeast strains: exploiting natural and artificial diversity. FEMS Microbiol Rev. 2014; 38: 947–995.

31. Wedral D, Shewfelt R, Frank J. The challenge of Brettanomyces in wine. LWT Food Sci and Technol. 2010; 43: 1474–1479.

